# Multi-factorial examination of amplicon sequencing workflows from sample preparation to bioinformatic analysis

**DOI:** 10.1101/2022.09.26.509576

**Authors:** Travis J. De Wolfe, Erik S. Wright

## Abstract

The development of sequencing technologies to evaluate bacterial microbiota composition has allowed new insights into the importance of microbial ecology. However, the variety of methodologies used among amplicon sequencing workflows leads to uncertainty about best practices as well as reproducibility and replicability among microbiome studies. Using a bacterial mock community composed of 37 soil isolates, we performed a comprehensive methodological evaluation of 540 workflows, each with a different combination of methodological factors spanning sample preparation to bioinformatic analysis to define sources of artifacts that affect sensitivity, specificity, and biases in the resulting compositional profiles. Of the 540 workflows examined, those using the V4-V4 primer set enabled the highest level of concordance between the original mock community and resulting microbiome sequence composition. Use of a high-fidelity polymerase, or a lower-fidelity polymerase with increased PCR elongation time limited chimera formation. Bioinformatic pipelines presented a trade-off between the fraction of distinct community members identified (sensitivity) and fraction of correct sequences (specificity). DADA2 and QIIME2 assembled V4-V4 reads amplified by Taq polymerase resulted in the highest specificity (100%), but only identified 52% of mock community members. Using mothur to assemble and denoise V4-V4 reads resulted in detection of 75% of mock community members among the resulting sequences, albeit with marginally lower specificity (99.5%). Optimization of microbiome workflows is critical for accuracy and to support reproducibility and replicability among microbiome studies. These aspects will help reveal the guiding principles of microbial ecology and impact the translation of microbiome research to human and environmental health.

## BACKGROUND

The use of next-generation sequencing in biological research laboratories has offered important insight into the role of microbial ecology to environmental and human health. Workflows to generate compositional microbiome data from a sample requires several laboratory-based sample preparation and computational steps. First, genomic DNA (gDNA) is extracted from the bacteria present in individual samples through physical or chemical lysing (1-3). From the gDNA, typically the 16S rRNA marker gene is amplified using polymerase chain reaction (PCR) and labeled with sample-specific index sequences (4, 5). The amplicons are then pooled and sequenced using a single- or paired-end sequencing approach. Finally, resulting reads are computationally demultiplexed using the sample-specific index sequences. With paired-end sequencing, two reads (R1 and R2) for each 16S rRNA template are produced by the instrument and are typically denoised by trimming and removing low-quality sequences before or after merging.

While techniques to study the microbiome have been adopted across many scientific fields, standardization of workflows to produce microbiome data has lagged, prompting researchers to develop in-house workflows (6, 7). Thus, methodological preferences at each step of a custom workflow (e.g. differing 16S rRNA primer sets, polymerases, PCR cycling conditions, and/or implementation of different bioinformatic software (8-10) make it difficult to identify methodological choices that maximize a workflow’s ability to correctly return distinct sequences (sensitivity) while limiting the proportion of incorrect sequences (specificity). Furthermore, in-house workflows oftentimes do not contain steps to mitigate biases nor common artifacts including reagent kit contaminants or cross-talk among index sequences (11, 12).

A study using data from the Human Microbiome Project and METAgenomics of the Human Intestinal Tract project illustrated this challenge when it failed to demonstrate a significant association between body mass index and taxonomic composition despite prior reports of associations in both mice and humans (13). In this study, the authors suggested the failure to replicate previous findings was likely due, in-part, to different methodological options used to prepare samples for sequencing. Another study focused on the microbial ecology of coral species found discrepant results from amplicon sequence data prepared by two different labs whose protocols differed only in DNA polymerase and sequencing platform (14). The authors emphasized caution when comparing and interpreting studies that combine amplicon sequence data across studies, even when only subtle differences exist between methodologies. Thus, replication among microbiome studies may benefit from the establishment of best practices for preparation and analysis of microbiome samples. This is also important to consider for the eventual adoption of microbiome-based clinical diagnostics which can be critically impacted by inaccurate workflows (8, 9, 15-19).

To circumvent inaccuracies as a result of different workflow methodologies among microbiome studies, microbial ecologists have made progress investigating the impact of individual library preparation factors on sequencing results (for review see (10)). Several reports have affirmed the significant impact of 16S rRNA primer selection on resulting microbiota composition, which sometimes depends on other workflow factors like the DNA extraction kit (2) or the sequencing platform used (20, 21). Others have noted the importance of pairing specific sequencing platforms with the appropriate bioinformatic pipelines for data analysis (22) or using specific polymerases with the appropriately determined number of PCR cycles to minimize artifacts (23). However, a comprehensive multi-factorial assessment of microbiome workflows that also considers bioinformatic processing of reads is lacking. In this study, we selected several factors, spanning sample preparation to bioinformatic processing, which we suspected *a priori* would be the most impactful to microbiome workflow accuracy. With these data we sought to identify a set of factors that may improve sensitivity and specificity of a microbiome study and contribute toward improved reproducibility and replicability in the microbiome field.

## RESULTS

Prior to bioinformatic denoising, we observed i7 and i5 sequence combinations among the raw reads that were not representative of sample index combinations intentionally included in the sequencing run. Reads corresponding to these unexpected index combinations are considered cross-talk artifacts and their occurrence sets the lower-detection limit for sample reads by providing a background of reads with unknown origin when samples are multiplexed for sequencing. We found an average cross-talk rate of 0.23% among samples indexed with the 1-step PCR approach and an average of 0.27% among samples indexed with the 2-step PCR approach (**Supplemental figure 1**). Consistent with previous studies that found similarly low rates of cross-talk (< 1%) (12), we found that i7 and i5 index reads had significantly lower mean Q-score among unexpected combinations as compared to expected in each sequencing run (1-step PCR i7 expected Q34.8 ± 0.8, unexpected Q28.6 ± 3.3, *t*(352.9) = 31.3, *p* < 0.001, *g* = 2.3; 1-step PCR i5 expected Q35.0 ± 0.1, unexpected Q31.3 ± 3.1, *t*(305.1) = 21.1, *p* < 0.001, *g* = 1.5; 2-step PCR i7 expected Q33.8 ± 2.5, unexpected Q29.1 ± 4.4, *t*(417.1) = 14.2, *p* < 0.001, *g* = 1.3; 2-step PCR i5 expected Q34.5 ± 1.3, unexpected Q30.8 ± 4.7, *t*(306.6) = 12.2, *p* < 0.001, *g* = 1.0). There was no apparent difference across sample preparation methods, underscoring a potential benefit of removing reads with low-quality index sequences generated by the Illumina sequencing platform.

### Quality of raw reads depends on the sequence read and its corresponding 16S rRNA primer set

Rarified sequence reads were evaluated prior to programmatic denoising and amplicon assembly to determine the effect of different library preparation factors on basic read characteristics, including string edit distance to the nearest mock community member and mean read Q-scores. We found that sequence read (R1 vs. R2) and the corresponding 16S rRNA regions (V4-V4 vs. V3-V4 vs. V4-V5) had the largest effect sizes regarding read characteristics. Additionally, R1 sequences had a lower edit distance to their nearest mock 16S rRNA sequence and had improved Q-scores over the same cumulative fraction of R2 sequences (Edit distance: *D* = 0.76, *p* < 0.001; Q-scores: *D* = 0.65, *p* < 0.001) (**Figure 1A**). After trimming to the high-quality region of each read, the average length remained longer for R1 than the same cumulative fraction of R2 reads (*D* = 0.64, *p* < 0.001). Considering the 16S rRNA amplification region, reads produced with the V4-V5 primer set had a higher average edit distance to the nearest mock 16S rRNA sequences (V3-V4 vs. V4-V5: *D* = 0.35, *p* = 6.8E-13; V4-V4 vs. V4-V5: *D* = 0.42, *p* < 0.001), the lowest average Q-score compared to the same cumulative fraction of reads produced from the V3-V4 or V4-V4 primer sets sequences (V3-V4 vs. V4-V5: *D* = 0.47, *p* < 0.001; V4-V4 vs. V4-V5: *D* = 0.59, *p* < 0.001), and the shortest average length after trimming (V3-V4 vs. V4-V5: *D* = 0.68, *p* < 0.001; V4-V4 vs. V4-V5: *D* = 0.82, *p* < 0.001) (**Figure 1B**). The reads produced with the V4-V4 primer set had a lower average edit distance to their nearest mock 16S rRNA sequence and the highest average Q-score compared the same cumulative fraction of other primer sets examined (Edit distance: V3-V4 vs. V4-V4 *D* = 0.37, *p =* 1.9E-14; Q-scores: V3-V4 vs. V4-V4 *D* = 0.37, *p =* 1.9E-14). Taken together, these results suggest the highest overall quality prior to downstream bioinformatic processing was among R1 reads and that those corresponding to the V4-V5 region had lowest quality compared to the other regions evaluated. Though we are unable to confirm this, it is possible the PCR conditions selected in our evaluation were not optimal for the V4-V5 primer-region, leading to lower overall read quality.

**Figure 1.**
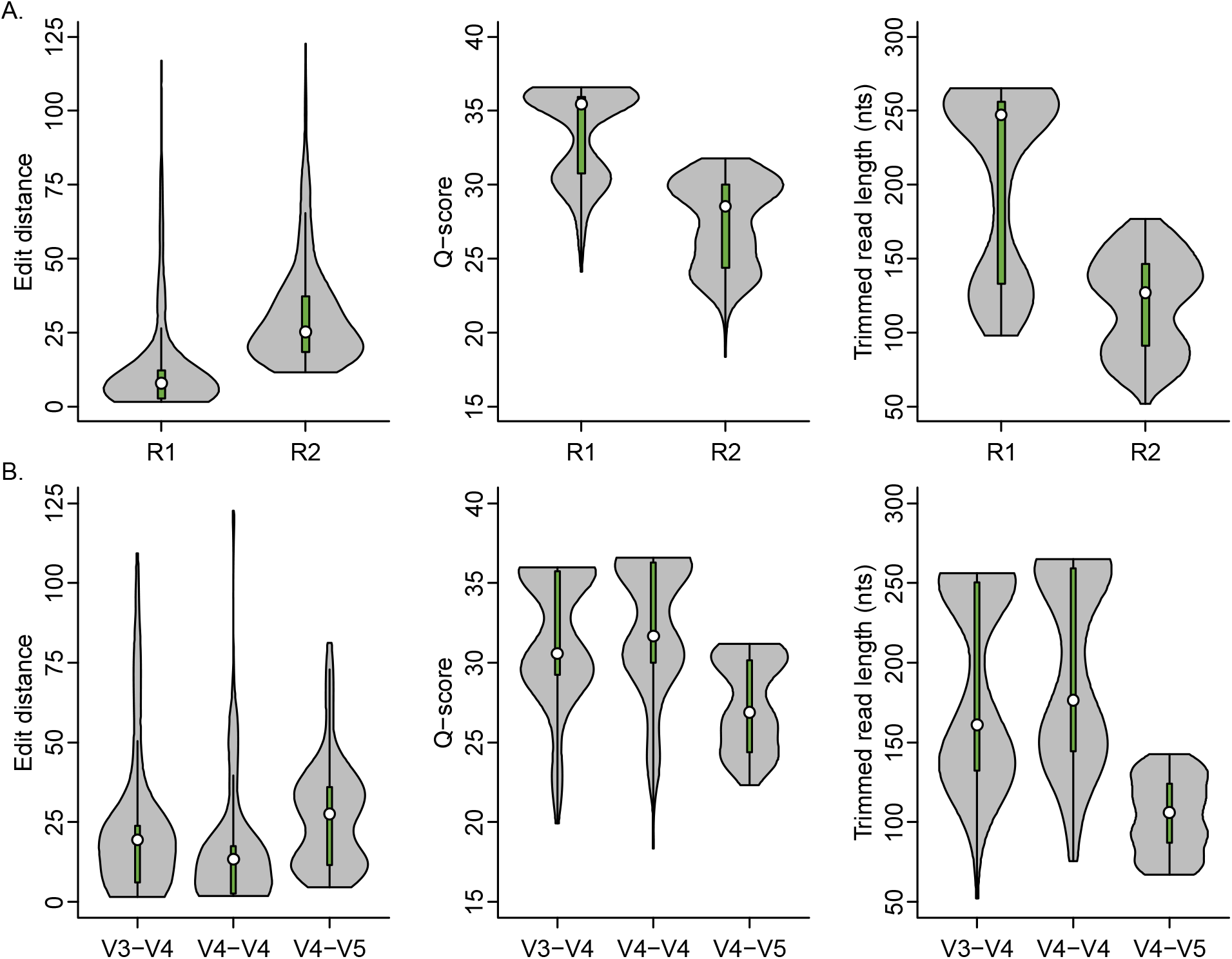
The sequence read and primer set influence edit distance and Q-score. A) R1 had consistently higher Q-scores than the same cumulative fraction of R2, likely limiting errors that would otherwise result in mismatches and allowing for greater overlap during amplicon assembly. B) Reads corresponding to the V4-V4 primer set had the lowest edit distance and highest Q-scores than the same cumulative fraction of reads corresponding to the V3-V4 and V4-V5 primer sets. Data shown represent kernel density estimates with green boxplots and white circles representing median values. Empirical cumulative distributions were significantly different (KS test *p* value < 0.001) among all factor levels compared within each plot.

### Exact matches to the mock community reveal a trade-off between specificity and sensitivity

Trade-offs between specificity and sensitivity are often observed when measuring the effect of experimental factors on accuracy. We estimated specificity as the fraction of resulting sequence variants that exactly matched the 16S rRNA sequence of the mock community members and sensitivity as the fraction of unique mock community members identified. We found that among raw reads, specificity and sensitivity were significantly impacted by primer set, polymerase, and the PCR indexing approach used. Overall, the specificity was low, indicating that a limited number of raw reads exactly matched the mock community sequences when taken immediately from the sequencer. Conversely, sensitivity was higher, indicating that despite many erroneous reads, the breadth of mock community members in the original sample was well represented. Raw reads corresponding with KAPA HiFi had higher specificity than both iTaq and SsoAdvanced (*F*(2, 306) = 136.2, *p* < 0.001, η^2^ = 0.23) **(Figure 2)**, and specificity was maximized when KAPA HiFi was paired with the V4-V4 primer set (*F*(4, 306) = 49.7, *p* < 0.001, η^2^ = 0.17) or 2-step PCR indexing (*F*(2, 306) = 50.8, *p* < 0.001, η^2^ = 0.09). Regardless of the protocol used, the V4-V4 primer set was associated with the highest specificity and sensitivity. Despite having the best overall sensitivity, the 1-step PCR indexing approach did not provide as high of specificity as 2-step PCR indexing (*F*(1, 306) = 85.2, *p* < 0.001, η^2^ = 0.07), especially when paired with the V4-V4 primer set. Finally, despite having lower specificity, iTaq and SsoAdvanced had significantly higher sensitivity than KAPA (*F*(2, 306) = 21.5, *p* < 0.001, η2 = 0.02), especially when paired with the V4-V4 primer set (*F*(4, 306) = 34.4, *p* < 0.001, η2 = 0.06).

**Figure 2.**
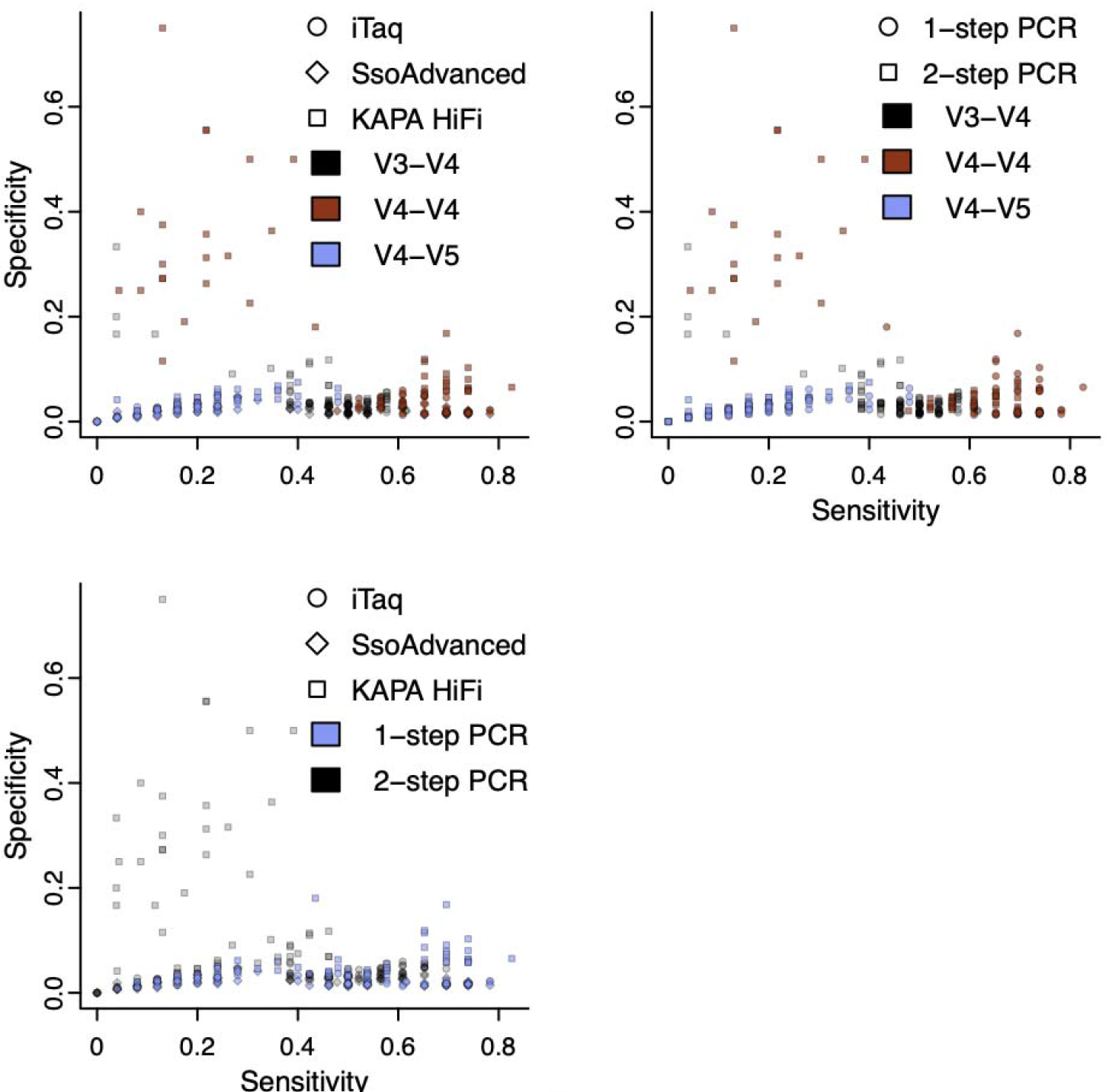
Sample preparation significantly impacts sensitivity and specificity of amplicon sequencing workflows prior to bioinformatic processing. The most sensitive workflow in this study could recall 83% of the original mock community members from the raw reads using the V4-V4 region of the 16S rRNA marker gene amplified with KAPA HiFi and 1-step PCR. The workflows with the highest specificity could only recall approximately 13% of the original mock community members from the raw reads using the V4-V4 region of the 16S rRNA marker gene amplified with KAPA HiFi and 2-step PCR.

After processing reads using each microbiome program, we found that specificity and sensitivity were impacted by primer set, polymerase, PCR indexing approach, and the specific bioinformatic processing program used. We observed lower specificity corresponding with the V3-V4 and V4-V5 primer sets (*F*(2, 464) = 171.3, *p* < 0.001, η^2^ = 0.24) **(Figure 3)**, and the highest sensitivity when samples were amplified with the V4-V4 primer set (*F*(2, 464) = 565.7, *p* < 0.001, η^2^ = 0.34). The specificity among different polymerase enzymes was lowest with KAPA HiFi and SsoAdvanced (*F*(2, 464) = 17.6, *p* = 4.5E-08, η^2^ = 0.02). When considering the PCR indexing approach, we observed significantly higher specificity (*F*(1, 464) = 97.4, *p* < 0.001, η^2^ = 0.07) and sensitivity with 1-step PCR (*F*(1, 464) = 391.1, *p* < 0.001, η^2^ = 0.12). Comparing bioinformatic processing programs, we found that mothur had significantly lower specificity than both DADA2 and QIIME2 due to a higher number of inexact matches among the representative OTUs (*F*(2, 464) = 13.6, *p* = 1.9E-06, ^2^ = 0.02). In contrast, the sensitivity of DADA2 and QIIME2 allowed for a maximal recall of approximately 52% of the original mock community sequences, which was significantly lower than the 75% maximally recalled using mothur (*F*(2, 464) = 233.9, *p* < 0.001, η^2^ = 0.14). Although data processed with mothur had lower specificity on average, it was just as high as that of DADA2 and QIIME2 when good quality data with fewer inexact matches was processed (mothur = 99.5% vs DADA2/QIIME2 = 100%).

**Figure 3.**
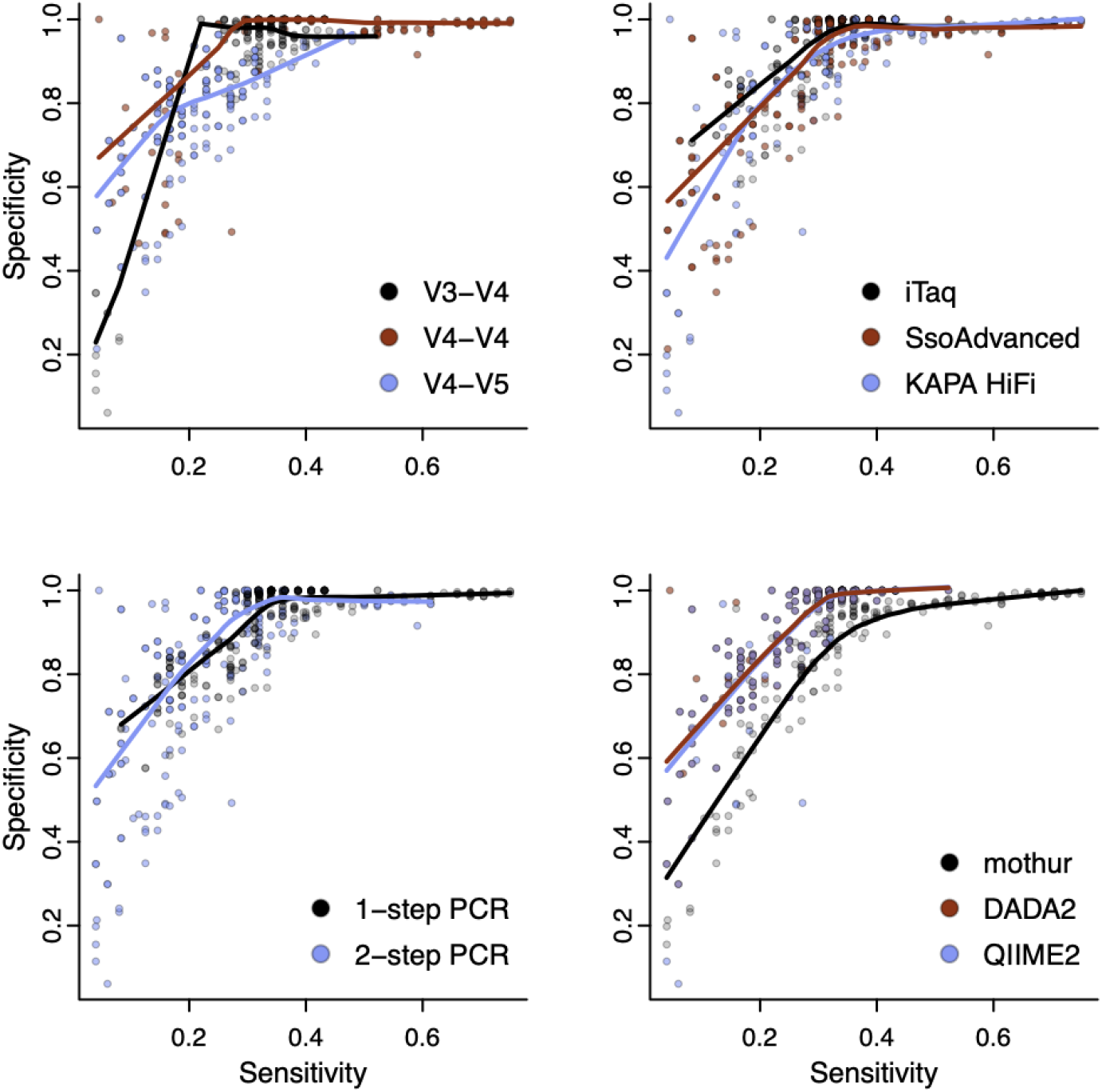
Trade-off between specificity and sensitivity depends on a combination of methodological factors. The most sensitive workflow in this study could recall 75% of the original mock community members using mothur to process reads from the V4-V4 region of the 16S rRNA marker gene amplified using 1-step PCR. The workflows with the highest specificity could only recall approximately 52% of the original mock community members using DADA2 or QIIME2. If high quality data was provided, the specificity of mothur processed data was as good as data processed with DADA2 or QIIME2.

### Primer-dimers and off-target amplification were uncommon among library preparation protocols

After classification and binning of exact matches, remaining inexact sequence variants were queried for primer sequences to identify primer-dimers. Among all samples, only 30 sequence variants (242 total reads, < 0.02%) were classified as primer-dimers. The successful elimination of primer-dimers was likely due to the size selectivity of the magnetic beads used for PCR library cleanup, allowing us to consider primer-dimers negligible artifacts in the present study. After primer-dimer classification, remaining inexact sequence variants were classified using IDTAXA to reveal three unique sequence variants (78 total reads) marked as potential off-target amplicons (**Supplemental table 1**). Subsequent BLAST analysis revealed that among them a single sequence variant matched to a 16S rRNA sequence in the database. The other two sequence variants did not match 16S rRNA sequences, did not have any significant hits (E-value < 0.01), and their nucleotide composition was of low complexity with homopolymeric DNA tracts. Therefore, these two sequence variants (77 total reads, < 0.01%) were likely sequencing failures or other sequencing artifacts. Like primer-dimers, the low prevalence of off-target amplicons allowed us to consider them negligible artifacts in this study.

### Chimeras are mitigated by increasing PCR elongation times

Chimeras were identified among inexact matches using a combination of program-specific chimera finding software (VSEARCH in mothur and *removeBimeraDenovo* in DADA2 and QIIME2), followed by *FindChimeras* from the DECIPHER package. The use of a second chimera finding method allowed us to identify chimeras not found by a microbiome bioinformatic processing pipeline (24). Overall, the fraction of original rarified sequences determined to be putatively chimeric was 0.03. There was a significant effect of elongation time on the fraction of chimeras detected (*F*(4, 1010) = 101.5, *p* < 0.001, η^2^ = 0.06), such that increasing elongation time reduced the average fraction of chimeric sequence reads from 0.04 (15 s) to 0.02 (180 s) over the range of elongation times tested. There was a significant interaction between elongation time and the primer set used (*F*(8, 1010) = 23.5, *p* < 0.001, η^2^ = 0.03), revealing that more chimeras were detected in reads corresponding to the V3-V4 primer set at elongation times ≤ 60 s as compared to the reads corresponding to the V4-V4 and V4-V5 primer sets (**Figure 4A)**. The polymerase used was found to be another important factor when considering elongation time and was consistent across primers (*F*(2, 1010) = 447, *p* < 0.001, η^2^ = 0.12), such that sequences amplified with KAPA HiFi generally had fewer chimeras detected regardless of elongation time (**Figure 4B**). Over the range of times tested, the fraction of chimeras detected with KAPA HiFi tended to decrease with increasing elongation time but was not significant, suggesting 30 s is sufficient to limit chimera formation with KAPA HiFi (*F*(4, 355) = 1.6, *p* = 0.18, η^2^ = 0.02).

**Figure 4.**
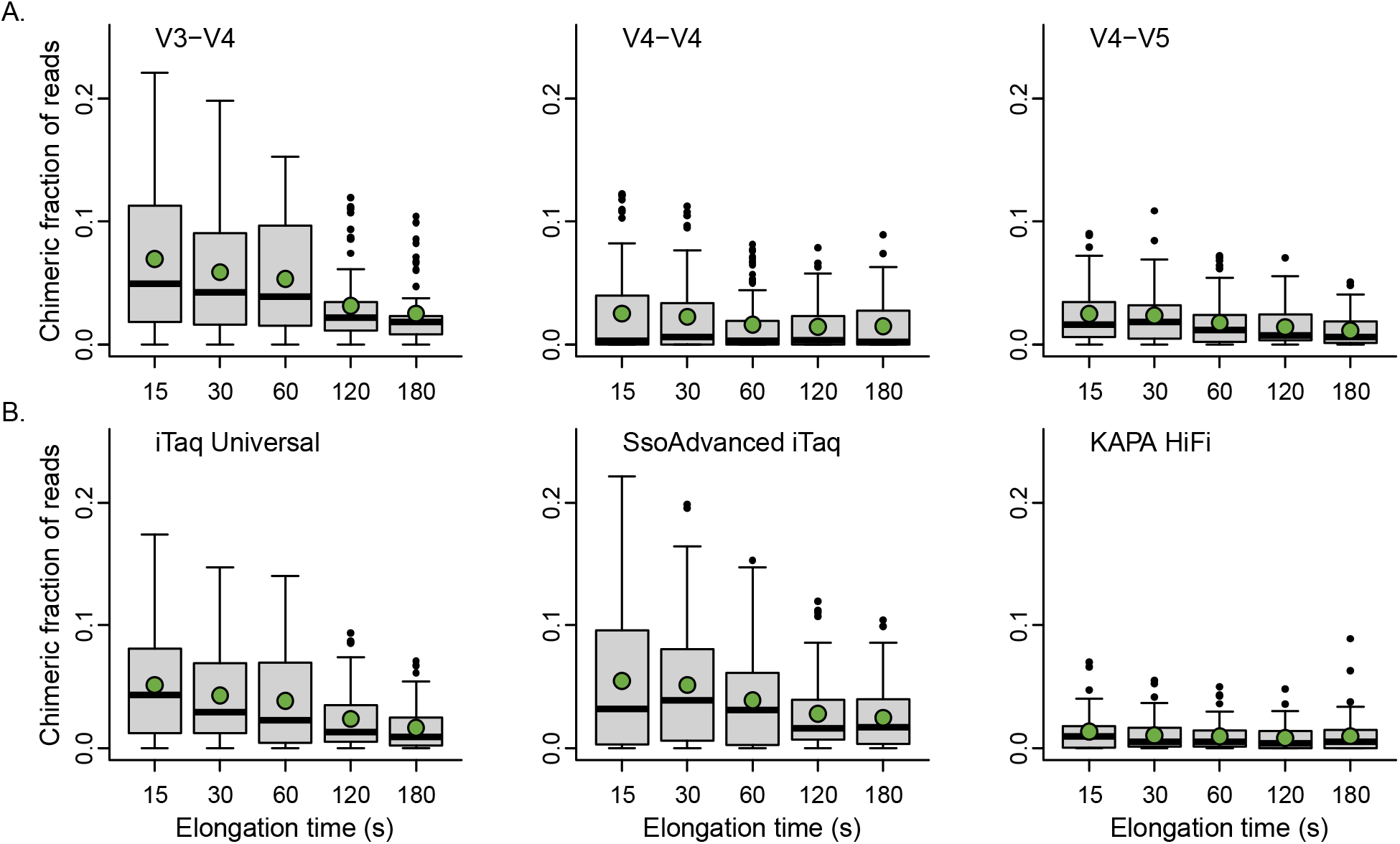
Increasing elongation time significantly reduced the fraction of chimeras that could be detected. A) Chimera detection remained highest for samples amplified with the V3-V4 primer set and B) lowest for samples amplified with KAPA HiFi polymerase. However, increasing elongation time with KAPA HiFi did not significantly reduce the fraction of chimeras detected. Data shown represent the putatively chimeric fraction of rarified sequence reads, with means signified with green circles and medians with wide black bars.

### Bioinformatic processing programs influence representation of rare sequence variants

Remaining inexact matches were classified as containing mismatches from the 16S rRNA sequences of mock community members or as contaminant sequences. We found that contaminants of low abundance were more prevalent after data processing by mothur than either QIIME2 or DADA2 (**Figure 5**). Evaluating the fate of the 359 contaminant reads that were removed by DADA2 and QIIME2 pipelines but not mothur revealed their omission with the *filterAndTrim* (67% of removed contaminants) or *dada* (33% of removed contaminants) functions. Upon examining the experimental origins of the contaminant reads, 71% were associated with specific sample subsets, with 42% of those originating from the negative control samples and the remaining 29% originating from samples prepared with the V3-V4 primer set. Those associated with the V3-V4 primer set are likely due to contamination introduced during commercial primer synthesis or handling within our laboratory. It is likely that the contaminants could be identified and removed using bioinformatic approaches such as with the decontam R package (25).

**Figure 5.**
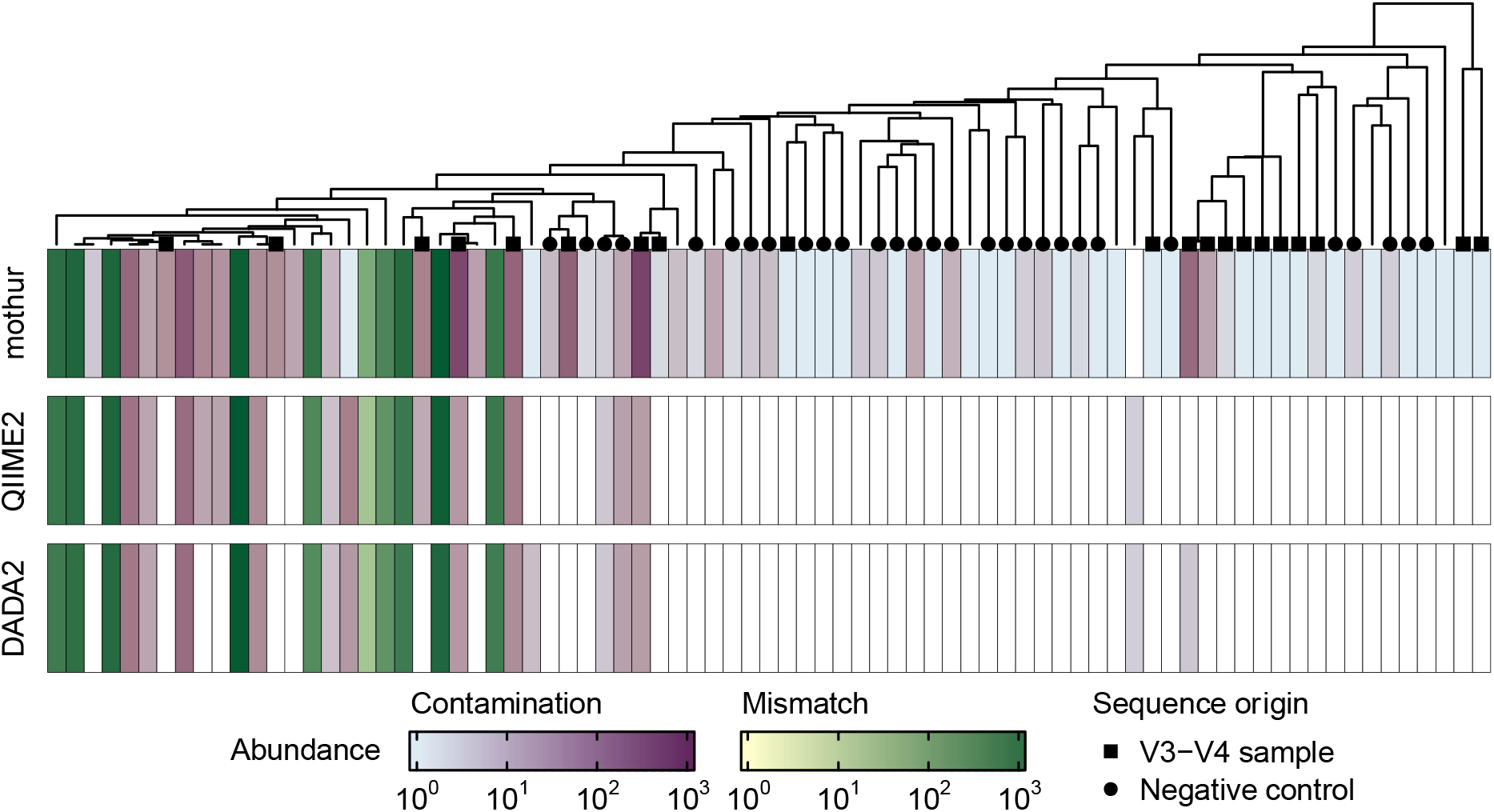
Detection of low abundance contaminants varied among bioinformatic processing programs. Contaminants were found primarily in mothur processed data and could largely be traced back to samples corresponding to the V3-V4 primer set or to negative controls. Data shown represent total read abundance for all inexact sequence variants collapsed into a consensus sequence within 15% distance. Consensus sequences were considered mismatches were if they clustered with a 16S rRNA sequence from the mock community. White heatmap cells indicate the consensus sequence was not represented among the corresponding data. The dendrogram represents a UPGMA tree constructed from the distances among consensus sequences.

### Methodological factors contribute to bias in observations of the community

Among microbiome studies, it is common to evaluate community composition through presence-absence or relative abundance of community members. Our evaluation of specificity and sensitivity of exact matches revealed that 1-step PCR with the V4-V4 primer set followed by bioinformatic processing with mothur best represented the expected presence of the original mock pool (precision). However, approximately 25% of the original mock community members were not identified in the best-case methodologies tested here. To incorporate the impact of methodological factors on relative abundance, we correlated the rank of each mock member’s approximate relative abundance in the original gDNA pool to the resulting relative abundance of exact matches among each methodology after bioinformatic processing. The correlations revealed that the polymerase, primer set, PCR indexing approach, and bioinformatic processing program can introduce significant bias into estimates of relative abundance. Using the V3-V4 or V4-V4 primer contributed a limited amount of bias relative to the V4-V5 primer set (*F*(2, 464) = 642.0, *p* < 0.001, η^2^ = 0.52) **(Figure 6)**. Evaluating different polymerases revealed KAPA HiFi contributed elevated bias compared to the others (*F*(2, 464) = 46.4, *p* < 0.001, η^2^ = 0.04) while the 1-step PCR indexing approach resulted in significantly higher correlation than 2-step PCR (*F*(1, 464) = 152.3, *p* < 0.001, η^2^ = 0.06). Bioinformatic processing with mothur best correlated with the relative abundance of mock members in the original DNA pool compared to DADA2 or QIIME2 (*F*(2, 464) = 6.2, *p* = 0.002, η^2^ = 0.005).

**Figure 6.**
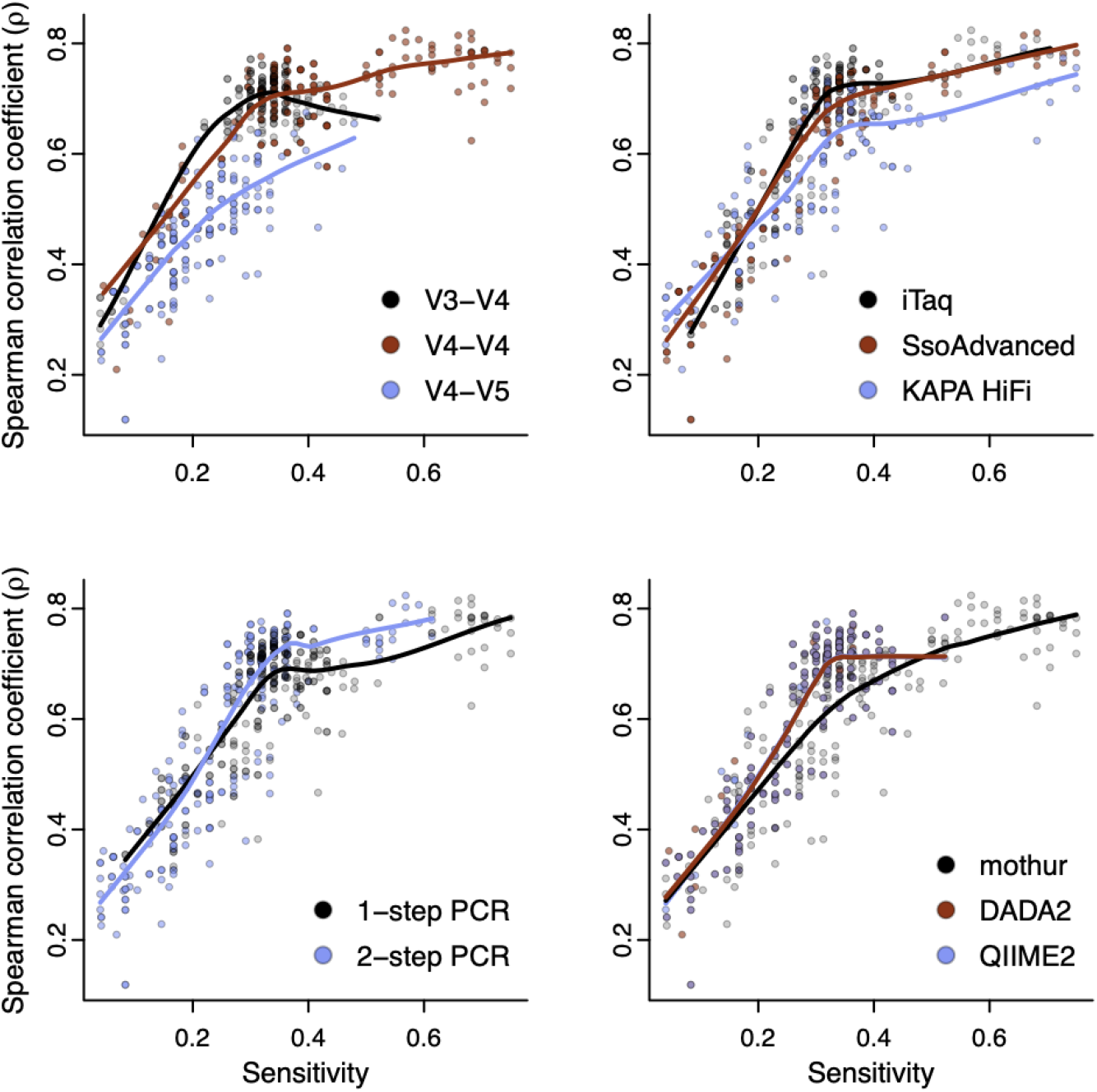
Compositional biases vary depending on the methodological factors used in each workflow. Correlation to the original mock community member estimated abundance was maximized when conducting 1-step PCR with the V4-V4 primer set and iTaq followed by denoising and read assembly with mothur. Data shown represent Spearman’s rank correlation between qPCR-derived approximate relative abundance of individuals originally included in the mock community to the relative abundance of exact sequence variants output from the bioinformatic processing programs.

## DISCUSSION

There is a growing appreciation for the sources and magnitude of errors and biases in microbiome data. This is largely a result of many methodological options available at each step of microbiome sample processing and bioinformatic analysis workflows and becomes critical when considering the degree of standardization required for translation of microbiome data to improving human and environmental health (9, 19, 26). Here, we reported a full-factorial evaluation of 540 microbiome workflows that included varying PCR cycling conditions, amplicon library preparation methods, and bioinformatic analyses. We found that all workflow factors introduce artifacts and biases to differing degrees which could impact downstream interpretation of microbiome composition. The only factor in which we did not detect meaningful differences was annealing temperature, but this may be due to the lack of mismatches between the primer sequences and 16S rRNA sequences of the our mock community members (27).

The 16S rRNA gene contains nine variable regions that differ in nucleotide sequence among bacterial phyla (**Supplemental figure 2**). Thus, ‘universal’ PCR primers are designed to amplify the chosen region and to maximize differentiation between bacterial groups in microbiome studies. Prior work suggests longer coverage of the 16S rRNA gene, such as using primers for V3-V4 or that combining sequences from different regions, allows for superior taxonomic resolution (28). This is supported by *in silico* studies that report optimal coverage of taxa in the SILVA database with the V3-V4 primer set and by verification among laboratory-based evaluations (29, 30). Like previous studies evaluating primers, we found that primers providing longer 16S rRNA coverage result in elevated error rates and reduced Q-scores among reads produced on the Illumina sequencing platform. These characteristics are critical to maximize the accurate merging of R1 and R2 into amplicons, as downstream denoising and assembly involves trimming reads to their higher quality region or culling reads if they are below a defined Q-score threshold (31). Until sequencing technologies are sufficiently accurate (32) we suggest selecting primer sets that provide shorter coverage of the 16S rRNA gene.

In addition to maximizing sequence read overlap, our assessment demonstrated additional advantages for using primers with shorter sequence coverage. Chimeras are PCR artifacts that can artificially inflate microbiome sample diversity and plague public taxonomy databases (33). We identified a higher prevalence of chimeras among sequence variants produced with the V3-V4 primer set compared to others when elongation time was greater than 60 s. Mapping nucleotide entropy over the length the 16S rRNA gene revealed that there is a conserved region with low entropy at the midpoint of the V3-V4 amplicon region (**Supplemental figure 2**). Prior investigations suggest these chimeric breakpoint locations are non-random and clustered around conserved sequence regions (34). Thus, it is likely that in combination with the read length produced from the V3-V4 primer set (approximately 432 nts), this conserved region promotes chimeras that can be detected with chimera finding software. Though the V4-V5 primer set also encompasses a conserved region, it is distally situated, likely reducing the probability of chimeras, or increasing the difficulty of their detection.

Regardless of the primer set used, increasing elongation time reduces the number of chimeras detected among the sequence variants when iTaq or SsoAdvanced polymerases are used. High-fidelity polymerases like KAPA HiFi have higher processivity, improved accuracy, and a lower error rate, allowing for continuous amplification and low dissociation from the DNA template. For these reasons, it is likely that each PCR cycle with KAPA HiFi resulted in complete synthesis, thus limiting the frequency of chimera formation even with only a 30 s elongation (35). Prior studies similarly highlighted the benefits of KAPA HiFi on chimera formation, including its lower rate of chimera formation relative to other polymerases which is independent of elongation time (23, 36, 37). It is critically important to mitigate chimera formation during PCR as bioinformatic algorithms likely cannot detect all chimeras (24).

We found that the presence of contaminant sequences depended upon which bioinformatic processing program was used. Specifically, low-abundance contaminants were primarily retained among data processed by mothur. Low-abundance contaminants were largely removed during the filter and trimming process, which screens for high quality sequences over a threshold sequence length, and during the process of inferring sample ASV composition. Many low-abundance mismatches and contaminants were represented by a single read after processing with mothur and was thus automatically removed by DADA2 and QIIME2. Further, some mismatch and contaminant reads from mothur processed data had lower Q-scores at specific positions along their length, suggesting that the user-definable criteria during filtering (e.g., a threshold for the maximum number of expected errors) also accounts for differences observed between programs. Recent work has found that the practice of removing rare sequences distorts the accurate compositional representation in a sample, making it more difficult to detect differences between treatments and disproportionately impacting samples with lower sequence depth (38). Further, this practice may be problematic for some research goals, particularly among studies investigating the rare biosphere, dietary profiles, or topology of trophic networks (39, 40). In these investigations it is preferred to identify those sequences that are not ubiquitous among samples and are distant to BLAST matches.

Using a mock community for our studies gave us the benefit of *a priori* knowledge of sequence representation, providing the confidence to identify contaminant sequences among mothur processed data that may have otherwise been considered part of the rare biosphere. Evaluating the factor-based origins of the contaminant reads, we found the V3-V4 primer set likely served as an inadvertent source of contamination due to sample handling or from the “kit-ome” (11). These contaminants can have a profound impact on samples with low biomass, where differences in types and abundances of contaminant DNA may drive differences in composition between samples (41-43). Thus, including negative controls in each sequencing run of a microbiome study is critical for determining background contamination and sequencing error rates. Furthermore, data from negative controls can be used to differentiate and remove contaminants (11, 44, 45)

We also identified contaminants in the negative control samples that could not be tied to a specific methodological factor. We believe these reads are a result of sample cross-talk, which is thought to be due to cluster overlap on the flow cell during Illumina sequencing (12, 46). In anticipation of evaluating cross-talk, we generated reads from every possible index combination, including those that were not used for a sample, allowing us to determine that cross-talk rates in our sequencing runs fell in line with what is typically observed in the absence of excess adapter (12, 47-49). Overall, the unexpected index combinations tended to have poor quality index reads and the corresponding quality was below Q30. It is important to note that without generating reads from every possible index combination, it would have been impossible to dismiss the impact of cross-talk in our study. Without mitigation by quality filtering index reads, or in the case that negative controls are not sequenced in each run, programs may unknowingly inflate sample diversity metrics through cross-talk. Thus, with the rapid and recent emergence of sequencing centers that are establishing microbiome pipelines, we repeat the call for cross-talk evaluation to become part of standard microbiome data analysis workflows (50).

Our findings underscored a trade-off between specificity and sensitivity a researcher must consider when designing a microbiome study. Specifically, we observed among the raw data that overall specificity was remarkably low, particularly for workflows with acceptable levels of sensitivity. The most sensitive workflow in our study could recall 83% of the original mock community members among the raw data, demonstrating the maximal sensitivity we would expect to observe after denoising. As this workflow employed KAPA HiFi to amplify the V4-V4 region, it is likely a reduced incorporation of chimeric reads along with the overall advantage of using the V4-V4 primer set which contributed to its accuracy.

An impactful finding of our study is that sample processing steps were found to interact with bioinformatic processing methods. We observed a significant global improvement in specificity after denoising and when assembling and denoising the reads with mothur, specificity was highest while also maximizing sensitivity and limiting compositional biases. We found that amplifying reads from the V4-V4 region using 1-step PCR with iTaq and ≥ 120 s elongation time or KAPA HiFi with a 30 s elongation time resulted in accurate sequences after denoising while using the V4-V5 primer set resulted in poor quality sequences. Lastly, using iTaq or SsoAdvanced with short elongation times (≤ 60 s) resulted in high rates of chimera formation that may have negative downstream impacts on sample diversity estimates. Recent studies have underscored the problematic nature of uncommon and spurious taxa among 16S rRNA data for confounding richness estimates and comparisons (51). This problem becomes compounded when attempting to use data at low taxonomic levels. The 16S rRNA gene is reported to have poor phylogenetic concordance and it is preferred to use whole genome sequences or multiple coding ribosomal gene sequences for species delineation instead (52). While the negative impacts of inaccurate workflows depend upon the downstream application, a small number of errors may still be correctly classified and result in the same conclusions. Thus, our aim is to evaluate workflow output without assuming downstream applications.

There are limitations to our study that are important for consideration. First, specific recommendations for microbiome workflows are difficult to make due to the limited phylogenetic diversity and structure of the mock community we used. Although it is one of the more diverse communities used in these types of studies, it does not recapitulate the possible diversity of bacteria among environmental samples (53). Furthermore, the mock community we used likely impacted the performance of the bioinformatic processing programs tested. If the parameters among each program were customized to our mock community we could expect alternative outcomes, however, we assumed that many microbiome researchers employ default standard operating procedures provided by the software developers (54). Therefore, we aimed to examine the data under these typical conditions. Another limitation is that our study represents a “simplistic” model in that our community is made using purified and pooled gDNA which excludes both sample specific characteristics such as presence of inhibitors as well as the impacts of other attributes including choice of DNA extraction methodology or starting template concentration.

## CONCLUSIONS

With a lack of standardized methodologies, reproducibility and replicability among microbiome studies remains problematic. In our evaluation of 540 different methodological combinations of sample preparation and bioinformatic analysis workflows, we identified important factors to improve specificity and sensitivity of a microbiome study while limiting biases. We believe these results underscore which parameters have greater influence over the accuracy of a microbiome study and provides a framework for optimizing workflows to a researcher’s needs. This work is important at the level of an individual study to determine guiding principles of microbial ecology but also toward the wider goal of addressing the reproducibility crisis that exists in the rapidly advancing microbiome field.

## METHODS

Purified bacterial isolates (n = 37) from soil with near full-length 16S rRNA gene sequences determined by Sanger sequencing (GenBank MN186620-MN186656) were inoculated on individual Luria-Bertani agar plates and incubated overnight (28 °C) (**Supplemental figure 3**). A single colony of each isolate was transferred to individual 0.2 mL thin-wall PCR tubes containing 100 µL sterile DNA grade water and sonicated at 50% amplitude for 60 s using a sonic dismembrator with cup horn attachment (55). Each suspension was subsequently centrifuged (60 s at 14,000 relative centrifugal force) and the resulting input supernatants containing gDNA were quantified using qPCR. The approximate 16S rRNA copy number of each isolate was determined using V4-V4 primers with 58 °C annealing temperature (**Table 1**). Next, gDNA from each isolate was pooled at equal volumes to serve as an uneven mock bacterial community of known presence and approximate abundance. Some divergent isolates were found to be polymorphic in specific regions of the 16S rRNA sequence while others were identical in a region. Thus, we expected that the 37 isolates corresponded to 27 unique sequence variants for the V3-V4 region, 23 sequence variants for the V4-V4 region, and 25 sequence variants for the V4-V5 region.

To compare the impact of library preparation factors, we performed all possible methodological combinations (180 combinations total) in duplicate for three polymerases, three alternative primer pairs, two indexing approaches (1-step or 2-step), five alternative elongation times, and two annealing temperature offsets (**Supplemental table 2**). PCR was performed using primers flanking either the V3-V4 or V4-V4 regions of the 16S rRNA gene or the top-scoring primer pair predicted by the DesignSignatures webtool (**Table 1**) (24, 56). The DesignSignatures primer set spans the V4-V5 region and was generated with the default parameters to maximize sensitivity and specificity to 6,482 genera in the Genome Taxonomy Database r89 bacterial and archaeal 16S rRNA database (57). For the 1-step PCR indexing approach, primers consisted of the 16S rRNA region-specific sequence, an eight nucleotide (nt) sequence index, and adapters specific to the Illumina MiSeq. All primer sets were subjected to gradient PCR to determine their empirical melting temperature (T_m_) with the pooled mock bacterial community gDNA as a template (**Supplemental table 3**).

**Table 1.**
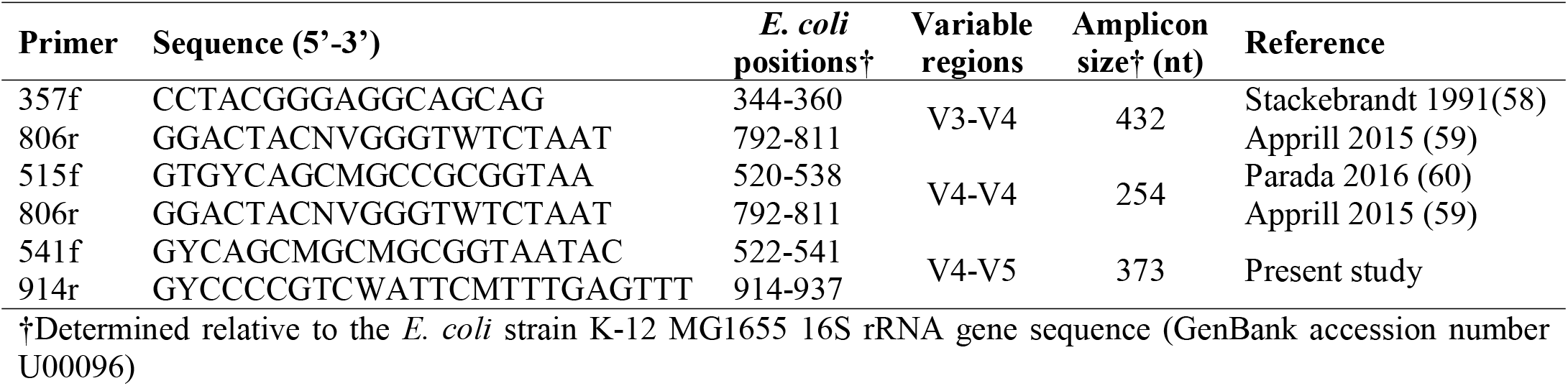
primers used in the multi-factorial study.

### PCR indexing

For 1-step PCR indexing, a total of 2 µL of each primer (10 µM) and 5 µL PCR pre-mix (iTaq Universal SYBR Green Supermix, SsoAdvanced Universal SYBR Green Supermix, or KAPA HiFi HotStart Real-time PCR Master Mix) were used for each reaction. A total of 1 µL of pooled gDNA was then added to each reaction. The cycling conditions on an Applied Biosystems StepOnePlus Real-Time PCR system began with an initial denaturation of 98 °C for 15 s followed by 40 cycles of annealing for 30 s, and elongation at 80 °C for 15, 30, 60, 120, or 180 s. The resulting indexed amplicons were pooled into a single library.

PCR cycling conditions for 2-step PCR indexing were as described above but with corresponding non-indexed primer pairs. Following the initial PCR, the resulting amplicons were diluted 1:50 and used as a template for five additional cycles of amplification to incorporate the corresponding indices used in the 1-step PCR (5, 61, 62). A negative DNA template control was included for all 2-step PCR combinations; however, a single indexed primer pair was used among all negative controls that shared the same non-indexed primer pair. The resulting indexed amplicons were pooled into a single library separate from the 1-step PCR library.

### Library clean-up and sequencing

An equal volume of each pooled amplicon library was independently cleaned and concentrated using Agencourt AMPure XP PCR purification beads according to manufacturer’s instructions. The two libraries (1-step and 2-step PCR) were subsequently spiked with 15% PhiX control DNA and sequenced in independent runs by the Center for Medicine and the Microbiome at the University of Pittsburgh on an Illumina MiSeq System with custom sequencing primers and a 2 × 300 bp MiSeq Reagent Kit v3 (**Supplemental table 4**). Sequences were demultiplexed and converted from base call files to FASTQ files using MiSeq Reporter software (v.2.6.2.3). FASTQ files for both index reads (i7 and i5) were generated by specifying the *CreateFastqForIndexReads* setting in the MiSeq Reporter.exe.config file. Raw reads are available from the Sequence Read Archive (BioProject PRJNA638427).

### Bioinformatic workflow

Raw index reads and associated Phred-scaled index read quality (Q) scores were generated for every possible i7 and i5 combination in each sequencing run, including those combinations that were not specifically assigned to a sample. These were evaluated for cross-talk by calculating the proportion of reads assigned to unexpected versus expected index combinations (63). Sample sequence reads (R1 and R2) were controlled for uneven sampling by reviewing the overall sequence read count distribution and rarified for consistency by randomly sampling sequence reads with replacement approximately at the first quartile of each sequencing run (1-step PCR: 4000 reads per sample; 2-step PCR: 1000 reads per sample) prior to subsequent analyses (64).

Two approaches were undertaken to assess the impact of library preparation factors on the raw data. First, a custom script was used to determine mean string edit distance to the nearest mock community member for reads of each sample and mean Q-scores (65, 66). Raw sequence reads were then trimmed based on Q-score using the *TrimDNA* function in DECIPHER v.2.25.3 and remaining reads were evaluated for mean sample read length (67). Second, raw sequence reads for each sample were aligned with the *pairwiseAlignment* function and merged into amplicons with a minimum overlap of 50 nt. Amplicons were then trimmed to the high-quality region with a minimum length of 200 nt using *TrimDNA* and clustered with the *Clusterize* function at a similarity threshold of 100%. The resulting clusters were matched to the mock community sequences with the *vcountPattern* function to determine the quantity of clusters that exactly matched to the mock community.

All microbiome pipelines recommend read trimming as a preliminary step. Since the corresponding amplicon size of the V4-V4 primer set is approximately 254 nt, adaptor read through occurred with the 2 × 300 bp MiSeq Reagent Kit and required trimming prior to subsequent processing. Next, trimming lengths to maximize read merging during downstream bioinformatic processing were pre-determined with a moving average (window size of 10) to the mean Q-scores along the original raw R1 and R2 sequences of each primer set. Trimming lengths were selected such that mean Q-score remained above Q30 or the minimum merged overlap ofR1 and R2 remained at least 30 nts. Raw sequence reads from each of the 180 workflows were then processed using the microbiome programs DADA2, QIIME2, and mothur, resulting in a final total of 540 different workflow combinations.

The microbiome program DADA2 v.1.14.0 was applied independently to raw sequence data from each sample with standard parameters defined in the pipeline tutorial (68). Briefly, sequence reads were filtered for a maximum of two expected errors in each read and trimmed according to pre-determined lengths based on moving average Q-score as described. For 1-step PCR, V3-V4 reads were trimmed to 295 and 189 nts, V4-V4 reads to 0 (no truncation) and 189 nts, and V4-V5 reads to 207 and 194 nts for R1 and R2, respectively. For 2-step PCR the V3-V4 reads were trimmed to 287 and 173 nts, V4-V4 to 0 and 186 nts, and V4-V5 to 187 and 214 nts. The DADA2 inference algorithm was then applied to the resulting sequence reads prior to merging and amplicon sequence variant (ASV) construction. In the rare case that *mergePairs* failed for a particular sample, only R1 was used in the DADA2 pipeline (20). Chimeric sequences were identified and removed from the ASV table using the DADA2 function *removeBimeraDenovo*.

The command-line QIIME 2 core v.2020.2 distribution was installed within a conda v.4.8.3 environment and was applied independently to raw sequence data from each sample with standard parameters defined in the QIIME2 “Atacama soil microbiome” and “Moving pictures” tutorials (69). Briefly, sequence reads were processed using the q2-dada2 plugin (DADA2 v.1.10.0 and R v.3.5.1) for trimming, merging, chimera removal, and ASV determination (68). In the rare case that *qiime dada2 denoise-paired* failed to merge reads due to the low quality of a particular sample, *qiime dada2 denoise-single* was implemented for only the R1 read in the QIIME2 pipeline.

Mothur v.1.43.0 was also applied independently to raw sequence data from each sample with standard parameters defined in the MiSeq standard operating procedures (70). Briefly, sequence reads were merged with a maximum of 8 homopolymers, 0 ambiguous bases, and a maximum amplicon length depending on the corresponding primer set (V3-V4: 450; V4-V4: 275; V4-V5: 400) (**Supplemental figure 2**). A customized reference alignment was constructed for each primer set with the small subunit rRNA (SSU) SILVA database (v.r132) and only those amplicons aligning to the appropriate regions were retained (71). Unique amplicons were filtered and those with up to two nucleotide differences were pre-clustered. Next, chimeras were removed from the pre-clustered amplicons using VSEARCH v.2.13.3 and remaining reads were clustered into OTUs at ≥ 97% similarity cutoff (72). The most abundant sequence variant in each cluster was selected as the OTU representative using *get*.*oturep*.

### Results assessment

For each analysis pipeline, tables containing variant counts were exported along with a FASTA file containing each representative sequence variant. Representatives were then assigned to a category: exact match, primer-dimer, chimera, off-target amplification, mismatched, or contamination. Briefly, sample-wise sequence variants were evaluated using a custom script that extracts sequences matching the mock community 16S rRNA sequences exactly (string edit distance of zero excluding length differences). The inexact matches were then examined for substrings containing a match to the sample’s respective primer sequence. Those which matched were removed as primer-dimers. Remaining sequence variants were then examined for chimeras using DECIPHER’s *FindChimeras* web tool with short-length sequence parameters (24). All inexact matches that were not identified as primer-dimers or chimeras were classified using IDTAXA with a modified SSU SILVA database (v.r132) (71, 73). Sequences that could not be classified (“unclassified_Root”) were presumed off-target amplification and subsequently evaluated for sequence similarity using the Basic Local Alignment Search Tool (BLAST) web-interface v.2.10.0+ using the non-redundant nucleotide sequence database (74). Matched targets with the lowest Expect (E) values to each inexact match query sequence were examined to confirm those likely to be off-target amplification. Query sequences that were closest to a non-16S rRNA sequence were considered off-target amplification. The remaining sequence variants (those matching a 16S rRNA sequence in BLAST) along with the sequence variants previously classified as 16S rRNA with IDTAXA were presumed to be contaminants or mismatched variants of mock community members. Inexact query sequence variants were defined as mismatches if they were within the same clade as a mock reference sequence when clustered using the average-linkage method and a 15% distance cutoff. Any inexact query sequences remaining were considered contamination.

After classifying sequence variants from all workflows, we calculated the relative abundance of exact matches (mock member output relative abundance = mock member read count / sum of read counts for all exact matches in the corresponding workflow), specificity (specificity = number of sequence reads from a workflow that exactly match mock sequence variants / total number of denoised sequence reads from the same workflow), and sensitivity (sensitivity = number of unique sequence variants from a workflow that exactly match mock sequence variants / total number of expected unique sequence variants for the same workflow) for each workflow. For example, if there are 1192 reads remaining for a workflow after denoising and 1141 of them exactly match a mock sequence variant, the specificity is 1141 / 1192 = 0.95. If the reads from this same workflow that used the V4-V5 primer set collapses into 10 unique sequence variants (ASVs or OTUs) that exactly match mock sequence variants, the sensitivity for the workflow is 10 / 25 = 0.4. Compositional bias was then determined by correlation between the qPCR estimated (input) relative abundance of individuals and the relative abundance of exact matches (output) with Spearman’s rank correlation coefficient (ρ) using the *cor* function in R. Workflows without output exact matches were omitted from the correlation.

### Statistical analyses

R v.4.0.3 was used for all statistical analyses (75). The impact of each methodological factor on raw R1 and R2 sequences was evaluated by calculating their respective proportion of variation explained (proportion of variation explained = factor sum of squares / total sum of squares). Differences between expected and unexpected index combinations for cross-talk were calculated using the *t*.*test* function and represented as mean ± standard deviation with effect size reported using the Hedges *g* (*g*) for independent groups (*g* = difference in group means / pooled and weighted standard deviation). Differences in distributions for the average edit distance, Q-score, and trimmed read length for the raw sequences were calculated using a two-sided Kolmogorov–Smirnov test (KS test) with the *ks*.*test* function in R and effect size (*D*) provided by the KS test statistic. Analysis of variation with consideration of second-order interaction effects was performed with the *aov* function and used to evaluate differences among means of sensitivity, specificity, and ρ for factors relating to categorically binned sequences. For all significant differences in factors, we computed the Tukey’s Honestly Significant Difference with *TukeyHSD* to assess the significance of differences between pairs of group means and eta squared η^2^) to compute the percentage of variance accounted for by each factor (η^2^ = sum of squares between factors / sum of squares total). Lines of best fit for scatter plots were smoothed with *lowess* in R. Differences among groups were considered statistically significant if the calculated *p* value was less than 0.05. Heatmaps were constructed with the R packages reshape2 v.1.4.4, circlize v.0.4.12, and ComplexHeatmap v.2.7.1 (76-78). Violin plots representing factor sample average were constructed using the R package vioplot v.0.3.4 (79).

## LIST OF ABBREVIATIONS

nt: nucleotide
R1: first sequence read
R2: second sequence read
gDNA: genomic DNA
PCR: polymerase chain reaction
bp: base pair
*Taq*: *Thermus aquaticus*
T_m_: melting temperature
OTU: operational taxonomic unit
ASV: amplicon sequence variant
DADA2: Divisive Amplicon Denoising Algorithm 2
QIIME2: Quantitative Insights Into Microbial Ecology 2
Q: Phred-scaled index read quality
KS test: Kolmogorov–Smirnov test
SSU: small subunit rRNA
BLAST: Basic Local Alignment Search Tool
E: Expect

## DECLARATIONS

### Ethics approval and consent to participate

Not applicable

### Consent for publication

Not applicable

### Availability of data and materials

The sequence reads generated and analyzed within this study are available on the National Center for Biotechnology Information Sequence Read Archive (BioProject PRJNA638427) https://www.ncbi.nlm.nih.gov/bioproject/PRJNA638427 *Competing interests:* The authors declare that they have no competing interests *Funding:* TJD was supported by the University of Pittsburgh Biomedical Informatics Training Program funded by the National Institutes of Health, National Library of Medicine [5T15LM007059-32] and the Michael Smith Foundation for Health Research Trainee award [RT-2020-04-64]. This study was supported by startup funding from the University of Pittsburgh to ESW.

## Authors contributions

TJD generated, analyzed, and interpreted all data within this study. ESW assisted in data generation, analysis, and interpretation and composition of the manuscript. All authors read and approved the final manuscript.

## Acknowledgements

We thank Mohammed Rafi Arefin (ORCID, 0000-0001-9227-9587), Andrew Beckley (ORCID: 0000-0003-2096-5208), Caroline Birer (ORCID: 0000-0001-7204-2013), Maria Bond, Lin Chou, Nicholas Cooley (ORCID: 0000-0002-6029-304X), Ann Donnelly (ORCID: 0000-0002-6296-5401), Genelle Healey (ORCID: 0000-0003-1418-2856), Nithya Narayanan, Nishant Panicker, and Krizia-Ivana Udquim for laboratory assistance and helpful conversation which improved the manuscript. Additionally, we thank Adam Fitch, Barbara Methe, and the Center for Microbiome and Medicine for their assistance with sequencing.

## REFERENCES

1. Douglas CA, Ivey KL, Papanicolas LE, Best KP, Muhlhausler BS, Rogers GB. DNA extraction approaches substantially influence the assessment of the human breast milk microbiome. Sci Rep. 2020;10(1):123.

2. Fouhy F, Clooney AG, Stanton C, Claesson MJ, Cotter PD. 16S rRNA gene sequencing of mock microbial populations-impact of DNA extraction method, primer choice and sequencing platform. BMC Microbiol. 2016;16(1):123.

3. Zhang D, Li W, Zhang S, Liu M, Gong H. Evaluation of the impact of DNA extraction methods on BAC bacterial community composition measured by denaturing gradient gel electrophoresis. Lett Appl Microbiol. 2011;53(1):44–9.

4. Kozich JJ, Westcott SL, Baxter NT, Highlander SK, Schloss PD. Development of a dual-index sequencing strategy and curation pipeline for analyzing amplicon sequence data on the MiSeq Illumina sequencing platform. Appl Environ Microbiol. 2013;79(17):5112–20.

5. Berry D, Ben Mahfoudh K, Wagner M, Loy A. Barcoded primers used in multiplex amplicon pyrosequencing bias amplification. Appl Environ Microbiol. 2011;77(21):7846–9.

6. Lebret K, Schroeder J, Balestreri C, Highfield A, Cummings D, Smyth T, et al. Choice of molecular barcode will affect species prevalence but not bacterial community composition. Mar Genomics. 2016;29:39–43.

7. Boers SA, Jansen R, Hays JP. Suddenly everyone is a microbiota specialist. Clin Microbiol Infect. 2016;22(7):581–2.

8. Bharti R, Grimm DG. Current challenges and best-practice protocols for microbiome analysis. Brief Bioinform. 2019.

9. Sinha R, Abnet CC, White O, Knight R, Huttenhower C. The microbiome quality control project: baseline study design and future directions. Genome Biol. 2015;16:276.

10. Pollock J, Glendinning L, Wisedchanwet T, Watson M. The Madness of Microbiome: Attempting To Find Consensus “Best Practice” for 16S Microbiome Studies. Appl Environ Microbiol. 2018;84(7).

11. Salter SJ, Cox MJ, Turek EM, Calus ST, Cookson WO, Moffatt MF, et al. Reagent and laboratory contamination can critically impact sequence-based microbiome analyses. BMC Biol. 2014;12:87.

12. Wright ES, Vetsigian KH. Quality filtering of Illumina index reads mitigates sample cross-talk. BMC Genomics. 2016;17(1):876.

13. Finucane MM, Sharpton TJ, Laurent TJ, Pollard KS. A taxonomic signature of obesity in the microbiome? Getting to the guts of the matter. PLoS One. 2014;9(1):e84689.

14. Epstein HE, Hernandez-Agreda A, Starko S, Baum JK, Vega Thurber R. Inconsistent Patterns of Microbial Diversity and Composition Between Highly Similar Sequencing Protocols: A Case Study With Reef-Building Corals. Front Microbiol. 2021;12:740932.

15. Schloss PD. Identifying and Overcoming Threats to Reproducibility, Replicability, Robustness, and Generalizability in Microbiome Research. mBio. 2018;9(3).

16. Amos GCA, Logan A, Anwar S, Fritzsche M, Mate R, Bleazard T, et al. Developing standards for the microbiome field. Microbiome. 2020;8(1):98.

17. Ravel J, Wommack KE. All hail reproducibility in microbiome research. Microbiome. 2014;2(1):8.

18. McLaren MR, Willis AD, Callahan BJ. Consistent and correctable bias in metagenomic sequencing experiments. Elife. 2019;8.

19. Sinha R, Abu-Ali G, Vogtmann E, Fodor AA, Ren B, Amir A, et al. Assessment of variation in microbial community amplicon sequencing by the Microbiome Quality Control (MBQC) project consortium. Nat Biotechnol. 2017;35(11):1077–86.

20. Tremblay J, Singh K, Fern A, Kirton ES, He S, Woyke T, et al. Primer and platform effects on 16S rRNA tag sequencing. Front Microbiol. 2015;6:771.

21. Claesson MJ, Wang Q, O’Sullivan O, Greene-Diniz R, Cole JR, Ross RP, et al. Comparison of two next-generation sequencing technologies for resolving highly complex microbiota composition using tandem variable 16S rRNA gene regions. Nucleic Acids Res. 2010;38(22):e200.

22. Allali I, Arnold JW, Roach J, Cadenas MB, Butz N, Hassan HM, et al. A comparison of sequencing platforms and bioinformatics pipelines for compositional analysis of the gut microbiome. BMC Microbiol. 2017;17(1):194.

23. Sze MA, Schloss PD. The Impact of DNA Polymerase and Number of Rounds of Amplification in PCR on 16S rRNA Gene Sequence Data. mSphere. 2019;4(3).

24. Wright ES, Yilmaz LS, Noguera DR. DECIPHER, a search-based approach to chimera identification for 16S rRNA sequences. Appl Environ Microbiol. 2012;78(3):717–25.

25. Davis NM, Proctor DM, Holmes SP, Relman DA, Callahan BJ. Simple statistical identification and removal of contaminant sequences in marker-gene and metagenomics data. Microbiome. 2018;6(1):226.

26. Lynch SV, Ng SC, Shanahan F, Tilg H. Translating the gut microbiome: ready for the clinic? Nat Rev Gastroenterol Hepatol. 2019;16(11):656–61.

27. Sipos R, Székely AJ, Palatinszky M, Révész S, Márialigeti K, Nikolausz M. Effect of primer mismatch, annealing temperature and PCR cycle number on 16S rRNA gene-targetting bacterial community analysis. FEMS Microbiol Ecol. 2007;60(2):341–50.

28. Fuks G, Elgart M, Amir A, Zeisel A, Turnbaugh PJ, Soen Y, et al. Combining 16S rRNA gene variable regions enables high-resolution microbial community profiling. Microbiome. 2018;6(1):17.

29. Klindworth A, Pruesse E, Schweer T, Peplies J, Quast C, Horn M, et al. Evaluation of general 16S ribosomal RNA gene PCR primers for classical and next-generation sequencing-based diversity studies. Nucleic Acids Res. 2013;41(1):e1.

30. Thijs S, Op De Beeck M, Beckers B, Truyens S, Stevens V, Van Hamme JD, et al. Comparative Evaluation of Four Bacteria-Specific Primer Pairs for 16S rRNA Gene Surveys. Front Microbiol. 2017;8:494.

31. Mohsen A, Park J, Chen YA, Kawashima H, Mizuguchi K. Impact of quality trimming on the efficiency of reads joining and diversity analysis of Illumina paired-end reads in the context of QIIME1 and QIIME2 microbiome analysis frameworks. BMC Bioinformatics. 2019;20(1):581.

32. Callahan BJ, Grinevich D, Thakur S, Balamotis MA, Yehezkel TB. Ultra-accurate Microbial Amplicon Sequencing Directly from Complex Samples with Synthetic Long Reads. bioRxiv. 2020:2020.07.07.192286.

33. Hugenholtz P, Huber T. Chimeric 16S rDNA sequences of diverse origin are accumulating in the public databases. Int J Syst Evol Microbiol. 2003;53(Pt 1):289–93.

34. Porazinska DL, Giblin-Davis RM, Sung W, Thomas WK. The nature and frequency of chimeras in eukaryotic metagenetic samples. J Nematol. 2012;44(1):18–25.

35. von Wintzingerode F, Göbel UB, Stackebrandt E. Determination of microbial diversity in environmental samples: pitfalls of PCR-based rRNA analysis. FEMS Microbiol Rev. 1997;21(3):213–29.

36. Ahn JH, Kim BY, Song J, Weon HY. Effects of PCR cycle number and DNA polymerase type on the 16S rRNA gene pyrosequencing analysis of bacterial communities. J Microbiol. 2012;50(6):1071–4.

37. Kurata S, Kanagawa T, Magariyama Y, Takatsu K, Yamada K, Yokomaku T, et al. Reevaluation and reduction of a PCR bias caused by reannealing of templates. Appl Environ Microbiol. 2004;70(12):7545–9.

38. Schloss PD. Removal of rare amplicon sequence variants from 16S rRNA gene sequence surveys biases the interpretation of community structure data. bioRxiv. 2020:2020.12.11.422279.

39. Sogin ML, Morrison HG, Huber JA, Mark Welch D, Huse SM, Neal PR, et al. Microbial diversity in the deep sea and the underexplored “rare biosphere”. Proc Natl Acad Sci U S A. 2006;103(32):12115–20.

40. Littleford-Colquhoun BL, Freeman PT, Sackett VI, Tulloss CV, McGarvey LM, Geremia C, et al. The precautionary principle and dietary DNA metabarcoding: commonly used abundance thresholds change ecological interpretation. Mol Ecol. 2022.

41. Gschwind R, Fournier T, Kennedy S, Tsatsaris V, Cordier AG, Barbut F, et al. Evidence for contamination as the origin for bacteria found in human placenta rather than a microbiota. PLoS One. 2020;15(8):e0237232.

42. Eisenhofer R, Minich JJ, Marotz C, Cooper A, Knight R, Weyrich LS. Contamination in Low Microbial Biomass Microbiome Studies: Issues and Recommendations. Trends Microbiol. 2019;27(2):105–17.

43. Jervis-Bardy J, Leong LE, Marri S, Smith RJ, Choo JM, Smith-Vaughan HC, et al. Deriving accurate microbiota profiles from human samples with low bacterial content through post-sequencing processing of Illumina MiSeq data. Microbiome. 2015;3:19.

44. Witzke M, Gullic A, Yang P, Bivens NJ, Adkins PRF, Ericsson AC. Influence of PCR cycle number on 16S rRNA gene amplicon sequencing of low biomass samples. J Microbiol Methods. 2020:106033.

45. de Goffau MC, Lager S, Salter SJ, Wagner J, Kronbichler A, Charnock-Jones DS, et al. Recognizing the reagent microbiome. Nat Microbiol. 2018;3(8):851–3.

46. Nelson MC, Morrison HG, Benjamino J, Grim SL, Graf J. Analysis, optimization and verification of Illumina-generated 16S rRNA gene amplicon surveys. PLoS One. 2014;9(4):e94249.

47. Costello M, Fleharty M, Abreu J, Farjoun Y, Ferriera S, Holmes L, et al. Characterization and remediation of sample index swaps by non-redundant dual indexing on massively parallel sequencing platforms. BMC Genomics. 2018;19(1):332.

48. van der Valk T, Vezzi F, Ormestad M, Dalén L, Guschanski K. Index hopping on the Illumina HiseqX platform and its consequences for ancient DNA studies. Mol Ecol Resour. 2019.

49. MacConaill LE, Burns RT, Nag A, Coleman HA, Slevin MK, Giorda K, et al. Unique, dual-indexed sequencing adapters with UMIs effectively eliminate index cross-talk and significantly improve sensitivity of massively parallel sequencing. BMC Genomics. 2018;19(1):30.

50. Martiny JBH, Whiteson KL, Bohannan BJM, David LA, Hynson NA, McFall-Ngai M, et al. The emergence of microbiome centres. Nat Microbiol. 2020;5(1):2–3.

51. Kumar MS, Slud EV, Hehnly C, Zhang L, Broach J, Irizarry RA, et al. Differential richness inference for 16S rRNA marker gene surveys. Genome Biol. 2022;23(1):166.

52. Hassler HB, Probert B, Moore C, Lawson E, Jackson RW, Russell BT, et al. Phylogenies of the 16S rRNA gene and its hypervariable regions lack concordance with core genome phylogenies. Microbiome. 2022;10(1):104.

53. Bokulich NA, Rideout JR, Mercurio WG, Shiffer A, Wolfe B, Maurice CF, et al. mockrobiota: a Public Resource for Microbiome Bioinformatics Benchmarking. mSystems. 2016;1(5).

54. Prodan A, Tremaroli V, Brolin H, Zwinderman AH, Nieuwdorp M, Levin E. Comparing bioinformatic pipelines for microbial 16S rRNA amplicon sequencing. PLoS One. 2020;15(1):e0227434.

55. Wright ES, Vetsigian KH. Inhibitory interactions promote frequent bistability among competing bacteria. Nat Commun. 2016;7:11274.

56. Wright ES, Vetsigian KH. DesignSignatures: a tool for designing primers that yields amplicons with distinct signatures. Bioinformatics. 2016;32(10):1565–7.

57. Parks DH, Chuvochina M, Waite DW, Rinke C, Skarshewski A, Chaumeil P-A, et al. A standardized bacterial taxonomy based on genome phylogeny substantially revises the tree of life. Nature Biotechnology. 2018;36:996.

58. Stackebrandt E, Goodfellow M. Nucleic acid techniques in bacterial systematics. Chichester ; New York: Wiley; 1991. xxix, 329 p. p.

59. A A, S M, R P, L W. Minor revision to V4 region SSU rRNA 806R gene primer greatly increases detection of SAR11 bacterioplankton. Aquatic Microbial Ecology. 2015;75(2):129–37.

60. Parada AE, Needham DM, Fuhrman JA. Every base matters: assessing small subunit rRNA primers for marine microbiomes with mock communities, time series and global field samples. Environ Microbiol. 2016;18(5):1403–14.

61. Gohl DM, Vangay P, Garbe J, MacLean A, Hauge A, Becker A, et al. Systematic improvement of amplicon marker gene methods for increased accuracy in microbiome studies. Nat Biotechnol. 2016;34(9):942–9.

62. Rausch P, Rühlemann M, Hermes BM, Doms S, Dagan T, Dierking K, et al. Comparative analysis of amplicon and metagenomic sequencing methods reveals key features in the evolution of animal metaorganisms. Microbiome. 2019;7(1):133.

63. Kircher M, Sawyer S, Meyer M. Double indexing overcomes inaccuracies in multiplex sequencing on the Illumina platform. Nucleic Acids Res. 2012;40(1):e3.

64. Hong J, Karaoz U, de Valpine P, Fithian W. To rarefy or not to rarefy: robustness and efficiency trade-offs of rarefying microbiome data. Bioinformatics. 2022.

65. Wright ES. DECIPHER: harnessing local sequence context to improve protein multiple sequence alignment. BMC Bioinformatics. 2015;16:322.

66. Pagès H, Aboyoun P, Gentleman R, DebRoy S. Biostrings: Efficient manipulation of biological strings. R package version 2.54.02019.

67. Wright E. Using DECIPHER v2.0 to Analyze Big Biological Sequence Data in R. The R Journal 2016. p. 352–9.

68. Callahan BJ, McMurdie PJ, Rosen MJ, Han AW, Johnson AJ, Holmes SP. DADA2: High-resolution sample inference from Illumina amplicon data. Nat Methods. 2016;13(7):581–3.

69. Bolyen E, Rideout JR, Dillon MR, Bokulich NA, Abnet CC, Al-Ghalith GA, et al. Reproducible, interactive, scalable and extensible microbiome data science using QIIME 2. Nat Biotechnol. 2019;37(8):852–7.

70. Schloss PD, Westcott SL, Ryabin T, Hall JR, Hartmann M, Hollister EB, et al. Introducing mothur: open-source, platform-independent, community-supported software for describing and comparing microbial communities. Appl Environ Microbiol. 2009;75(23):7537–41.

71. Quast C, Pruesse E, Yilmaz P, Gerken J, Schweer T, Yarza P, et al. The SILVA ribosomal RNA gene database project: improved data processing and web-based tools. Nucleic Acids Res. 2013;41(Database issue):D590–6.

72. Rognes T, Flouri T, Nichols B, Quince C, Mahé F. VSEARCH: a versatile open source tool for metagenomics. PeerJ. 2016;4:e2584.

73. Murali A, Bhargava A, Wright ES. IDTAXA: a novel approach for accurate taxonomic classification of microbiome sequences. Microbiome. 2018;6(1):140.

74. Altschul SF, Gish W, Miller W, Myers EW, Lipman DJ. Basic local alignment search tool. J Mol Biol. 1990;215(3):403–10.

75. Team RC. R: A Language and Environment for Statistical Computing. 3.6.3 ed2020.

76. Gu Z, Eils R, Schlesner M. Complex heatmaps reveal patterns and correlations in multidimensional genomic data. Bioinformatics. 2016;32(18):2847–9.

77. Gu Z, Gu L, Eils R, Schlesner M, Brors B. circlize Implements and enhances circular visualization in R. Bioinformatics. 2014;30(19):2811–2.

78. Wickham H. Reshaping Data with the reshape Package. 2007. 2007;21(12):20.

79. Adler D, Kelly ST. vioplot: violin plot. R package version 0.3.4 2019 [Available from: https://github.com/TomKellyGenetics/vioplot.

