## Supplemental files for "Multi-factorial examination of amplicon sequencing workflows from sample preparation to bioinformatic analysis"

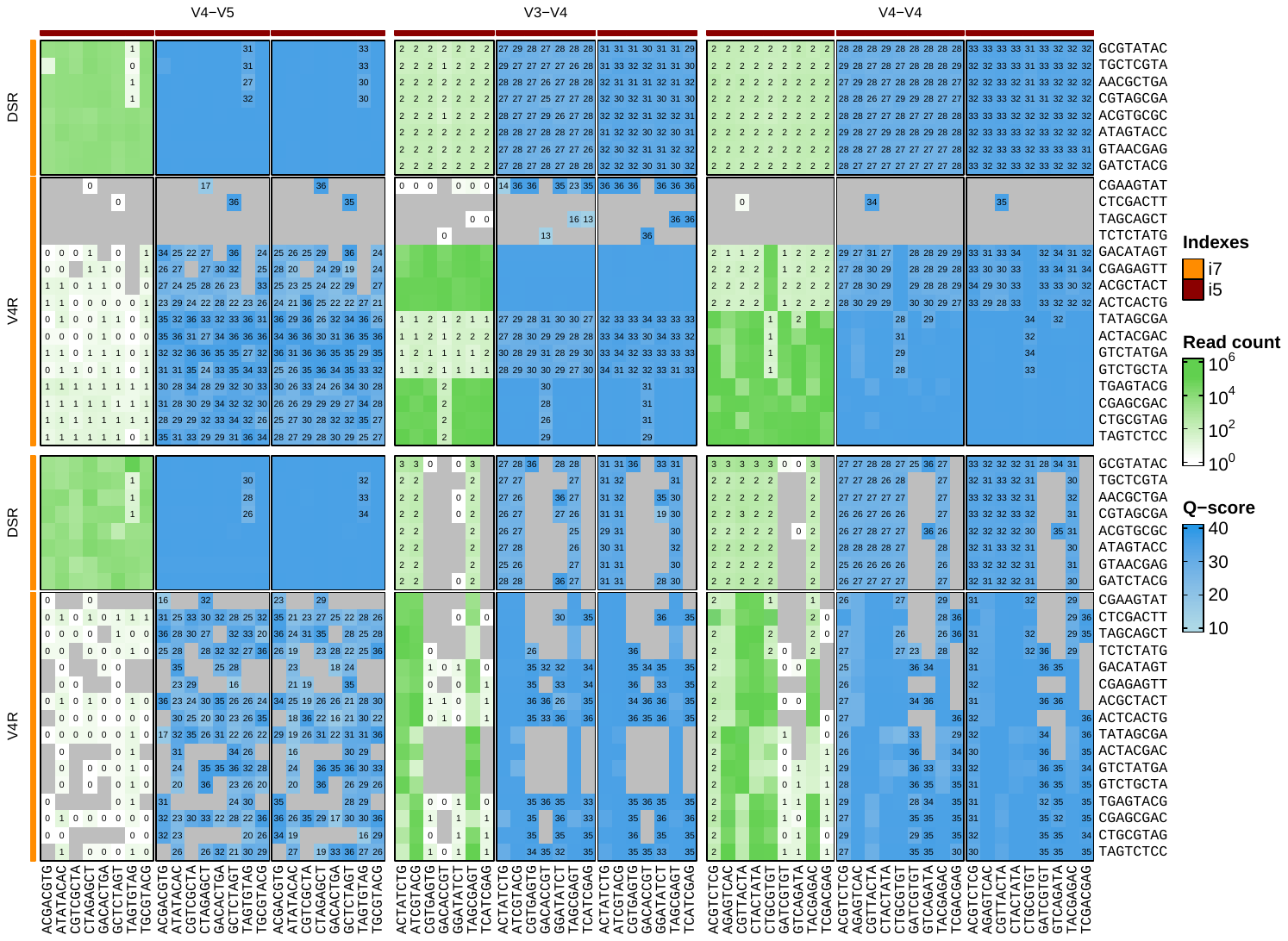

**Supplemental figure 1.** Index read count and Q-scores for expected or unexpected index combinations found among raw sequences. Heatmap cells containing a read count or Q-score for the respective index combination indicates the combination was not included as a sample in the sequencing run (unexpected, false positive). Grey heatmap cells indicate the index combination was not included as a sample in the sequencing run and it was not found in the resulting sequence data (true negative).

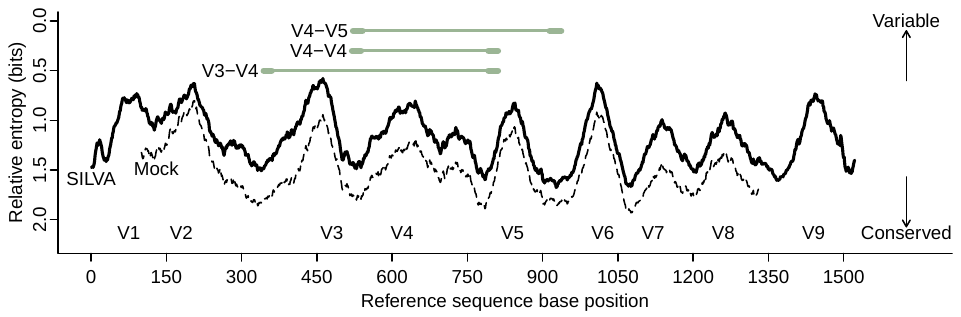

**Supplemental figure 2**. Entropy along the 16S rRNA reveals variable regions (V1 to V9) for inferring phylogenetic relationships. The primer set used in a 16S rRNA survey defines the variable region(s) that will be amplified during the PCR step of library preparation. Relative entropy values were determined from the SILVA r138 database and the mock bacterial community members used in this study. Base positions are numbered according to the *Escherichia coli* K-12 MG1655 16S rRNA positive-strand (GenBank accession number U00096). Note that the y-axis is reversed so variable regions display ‘high’ entropy on the plot.

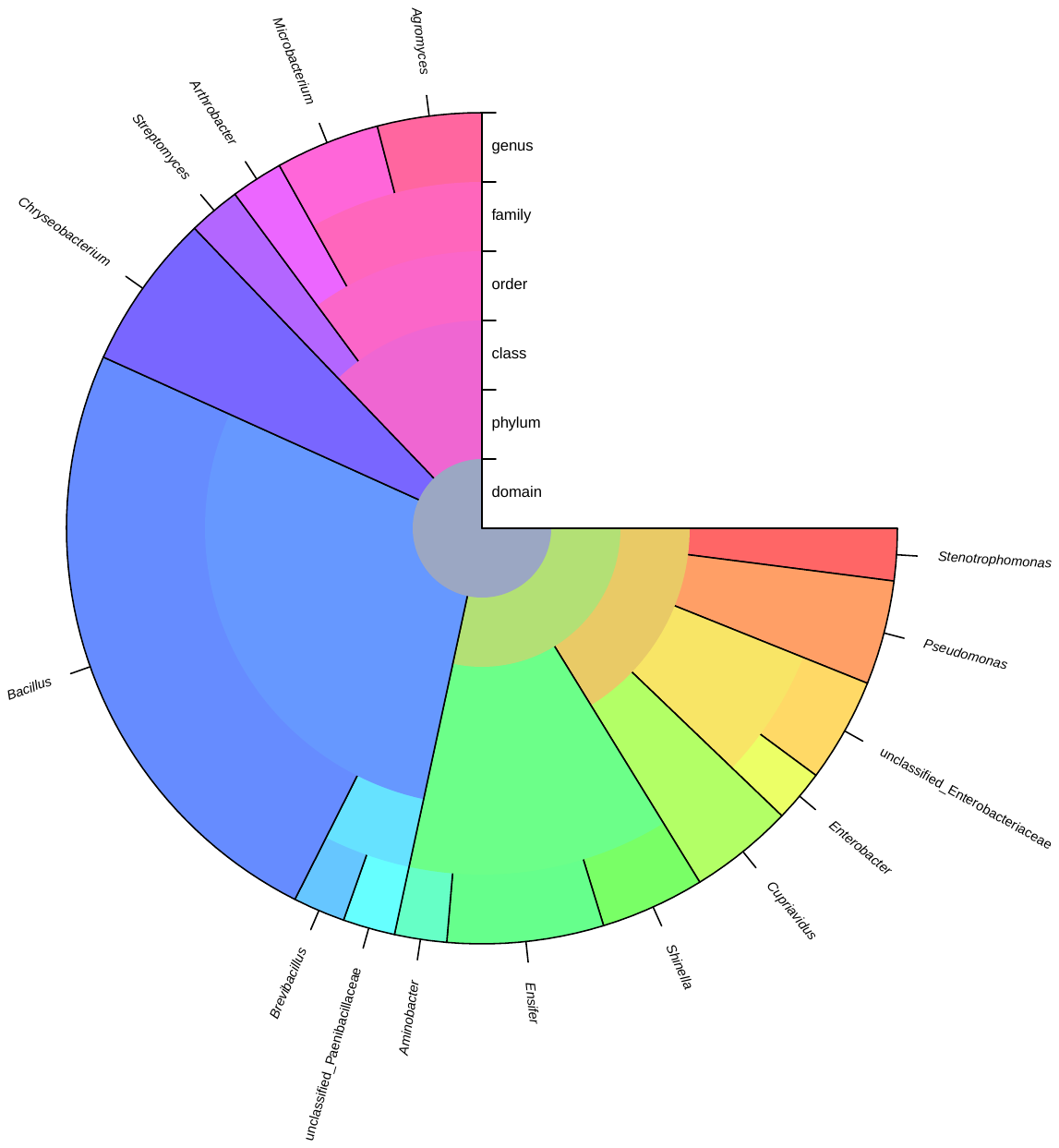

**Supplemental figure 3**. A total of 37 unique bacterial isolates were cultivated from soil and sequenced to construct a phylogenetic- and taxonomically diverse mock community. Taxonomy was assigned by IDTAXA with the SILVA r138 database.

| **Supplemental table 1.** BLAST hits for potential off-target amplicon | | | | | | |
| --- | --- | --- | --- | --- | --- | --- |
| **Hit accession number** | **Percent identity** | **E-value** | **Bit score** | **Description** | **Off-target amplicon*** | **Read count** |
| LN591134.1 | 79.03 | 0.066 | 50.9 | *Cyprinus carpio* | Y | 49 |
| LN591134.1 | 79.03 | 0.066 | 50.9 | *Cyprinus carpio* | Y | 28 |
| KR849800.1 | 100.00 | 5.73E-123 | 452.0 | Uncultured bacterium | N | 1 |
| *Inexact matches that could not be classified with IDTAXA were evaluated for sequence similarity using BLAST v.2.10.0+. Sequence hits not matching to 16S rRNA were classified as off-target amplification | | | | | | |

| **Supplemental table 2. Methodological factors evaluated in the present study** | | | | | |
| --- | --- | --- | --- | --- | --- |
| **Indexing approaches** | **16S rRNA regions** | **Polymerases** | **Elongation times** | **Annealing temperatures** | **Program** |
| 1-step PCR | V3-V4 | iTaq (Bio-Rad; 1725121) | 15 | T_m_ | DADA2 |
| 2-step PCR | V4-V4 | SsoAdvanced (Bio-Rad; 1725270) | 30 | 5 °C below T_m_ | QIIME2 |
|  | V4-V5 | KAPA HiFi (Kapa Biosystems; KK2702) | 60 |  | mothur |
|  |  |  | 120 |  |  |
|  |  |  | 180 |  |  |

| **Supplemental table 3. Primer-wise annealing temperatures used in the present study** | | | | |
| --- | --- | --- | --- | --- |
| **Primer** | **Polymerase** | **Indexing approach** | **Annealing temperature** | **Annealing temperature (°C)** |
| V3-V4 | iTaq | 1-step PCR | 5 °C below T_m_ | 53 |
| V3-V4 | SsoAdvanced | 1-step PCR | 5 °C below T_m_ | 53 |
| V3-V4 | KAPA | 1-step PCR | 5 °C below T_m_ | 62 |
| V3-V4 | iTaq | 1-step PCR | T_m_ | 58 |
| V3-V4 | SsoAdvanced | 1-step PCR | T_m_ | 58 |
| V3-V4 | KAPA | 1-step PCR | T_m_ | 67 |
| V4-V4 | iTaq | 1-step PCR | 5 °C below T_m_ | 53 |
| V4-V4 | SsoAdvanced | 1-step PCR | 5 °C below T_m_ | 53 |
| V4-V4 | KAPA | 1-step PCR | 5 °C below T_m_ | 62 |
| V4-V4 | iTaq | 1-step PCR | T_m_ | 58 |
| V4-V4 | SsoAdvanced | 1-step PCR | T_m_ | 58 |
| V4-V4 | KAPA | 1-step PCR | T_m_ | 67 |
| V4-V5 | iTaq | 1-step PCR | 5 °C below T_m_ | 53 |
| V4-V5 | SsoAdvanced | 1-step PCR | 5 °C below T_m_ | 53 |
| V4-V5 | KAPA | 1-step PCR | 5 °C below T_m_ | 62 |
| V4-V5 | iTaq | 1-step PCR | T_m_ | 58 |
| V4-V5 | SsoAdvanced | 1-step PCR | T_m_ | 58 |
| V4-V5 | KAPA | 1-step PCR | T_m_ | 67 |
| V3-V4 | iTaq | 2-step PCR | 5 °C below T_m_ | 53 |
| V3-V4 | SsoAdvanced | 2-step PCR | 5 °C below T_m_ | 53 |
| V3-V4 | KAPA | 2-step PCR | 5 °C below T_m_ | 62 |
| V3-V4 | iTaq | 2-step PCR | T_m_ | 58 |
| V3-V4 | SsoAdvanced | 2-step PCR | T_m_ | 58 |
| V3-V4 | KAPA | 2-step PCR | T_m_ | 67 |
| V4-V4 | iTaq | 2-step PCR | 5 °C below T_m_ | 53 |
| V4-V4 | SsoAdvanced | 2-step PCR | 5 °C below T_m_ | 53 |
| V4-V4 | KAPA | 2-step PCR | 5 °C below T_m_ | 62 |
| V4-V4 | iTaq | 2-step PCR | T_m_ | 58 |
| V4-V4 | SsoAdvanced | 2-step PCR | T_m_ | 58 |
| V4-V4 | KAPA | 2-step PCR | T_m_ | 67 |
| V4-V5 | iTaq | 2-step PCR | 5 °C below T_m_ | 53 |
| V4-V5 | SsoAdvanced | 2-step PCR | 5 °C below T_m_ | 53 |
| V4-V5 | KAPA | 2-step PCR | 5 °C below T_m_ | 62 |
| V4-V5 | iTaq | 2-step PCR | T_m_ | 58 |
| V4-V5 | SsoAdvanced | 2-step PCR | T_m_ | 58 |
| V4-V5 | KAPA | 2-step PCR | T_m_ | 67 |

| **Supplemental table 4. Custom sequencing primers** | |
| --- | --- |
| **Name** | **Sequence** |
| V3_R1 | TATGGTAATTGGCCTACGGGAGGCAGCAG |
| V4_R2 | AGTCAGTCAGCCGGACTACNVGGGTWTCTAAT |
| V34_V4_index | ATTAGAWACCCBNGTAGTCCGGCTGACTGACT |
| V4_R1 | TATGGTAATTGTGTGYCAGCMGCCGCGGTAA |
| V45_R1 | TATGGTAATTAAGYCAGCMGCMGCGGTAATAC |
| V45_R2 | AGTCAGTCAGAAGYCCCCGTCWATTCMTTTGAGTTT |
| V45_index | AAACTCAAAKGAATWGACGGGGRCTTCTGACTGACT |
